# Advanced Research Infrastructure for Experimentation in genomicS (ARIES): a lustrum of Galaxy experience

**DOI:** 10.1101/2020.05.14.095901

**Authors:** Arnold Knijn, Valeria Michelacci, Massimiliano Orsini, Stefano Morabito

## Abstract

Background: With the introduction of Next Generation Sequencing (NGS) and Whole-Genome Sequencing (WGS) in microbiology and molecular epidemiology, the development of an information system for the collection of genomic and epidemiological data and subsequent transparent and reproducible data analysis became indispensable. Further requirements for the system included accessibility and ease of use by bioinformatics as well as command line profane scientists.

Findings: The ARIES (Advanced Research Infrastructure for Experimentation in genomicS, https://aries.iss.it) platform has been implemented in 2015 as an instance of the Galaxy framework specific for use of WGS in molecular epidemiology. Here, the experience with ARIES is reported. Conclusions: During its five years existence, ARIES has grown into a well-established reality not only as a web service but as well as a workflow engine for the Integrated Rapid Infectious Disease Analysis (IRIDA) platform. In fact, an environment has been created with the implementation of complex bioinformatic tools in an easy-to-use context allowing scientists to concentrate on what to do instead of how to do it.

## Findings

### Background

The constant evolution of sequencing techniques and the parallel development of bioinformatics has opened new perspectives in public health microbiology [1,2,3]. Advances in molecular epidemiology include new insights in microbial typing, geographical as well as temporal diffusion of genomic data. This is particularly true in the present phase, during the change in bacterial typing paradigm, moving from Pulsed-field Gel Electrophoresis (PFGE) to Next Generation Sequencing (NGS)-based whole genome and core genome MultiLocus Sequence Typing (wg/cgMLST) [4]. In spite of the increasing usage of NGS to produce microbial typing data, still no golden standard has been defined for the analysis and comparison of the data produced with different available sequencing techniques [5,6,7] and the many bioinformatic solutions developed, which make use of the allelic or single nucleotide polymorphisms differences to come to a strain signature for phylogenetic analyses [1,8,9,10]. Thus, the demand is posed for the development of solutions able to provide reproducible analyses, equally accessible to scientists with and without specific informatics expertise. For this, the Galaxy Project offers an exquisite platform: an open-source web-based framework supported by a strong community delivering high-quality tutorials and broad integration of available bioinformatic tools [12]. The ARIES (Advanced Research Infrastructure for Experimentation in genomicS, https://aries.iss.it) platform has been implemented in 2015 as a medium-small instance of the Galaxy framework specific for use of Whole-Genome Sequencing (WGS) in molecular microbiology and epidemiology. Here a description of five years of experience with the platform is given.

### European Union context

The Directive 2003/99/EC of the European Parliament and of the Council [13] was emanated with the purpose to ensure that zoonoses, zoonotic agents and related antimicrobial resistance are properly monitored and that food-borne outbreaks receive proper epidemiological investigation. In this framework, the “European Union Reference Laboratory for *Escherichia coli*, including Verotoxigenic *E. coli*” (EURL VTEC) was assigned following a public call in 2006 to Microbiological Food Safety and Food-borne Diseases Unit of the Department of Food Safety, Nutrition and Veterinary Public Health of the Istituto Superiore di Sanità (ISS), Rome, Italy. The European Member States on their part have designated National Reference Laboratories (NRLs) that interact on one hand with the EURL and on the other with their local official food control laboratories. The responsibilities and tasks of EURLs (Regulation (EU) 2017/625, Art. 94) [14] are amongst others:

- to contribute to the improvement and harmonisation of methods of analysis, test or diagnosis;
- to provide NRLs with analytical reference methods;
- to organise comparative testing (Proficiency Tests);
- to conduct training courses for NRLs;
- to provide scientific and technical assistance to the European Commission;
- to assist actively in the diagnosis of outbreaks in Member States.

Furthermore, a EURLs Working Group on WGS was instituted in 2017 by the European Commission with the aim to promote the use of NGS and WGS across the EURLs’ networks and build this capacity within the EU. Also, liaison was to be ensured between the work of the EURLs and the work of the European Food Safety Authority (EFSA) and the European Centre for Disease Prevention and Control (ECDC) on the basis of a WGS mandate sent to them by the European Commission. The Working group is made up of eight EURLs and coordinated by the EURL VTEC.

### ARIES goals and evolution

In this context and following an early adoption of WGS for the molecular epidemiology of *E. coli* infections, the EURL VTEC contacted the Informatics Service of the ISS in 2014 in order to plan the provisioning of a platform that would allow the researchers working at the EURL VTEC, at a first instance, for simple access to genomic and molecular epidemiology bioinformatic tools. In April of the same year a pilot standalone installation of the Galaxy framework was performed (Figure 1). Extensive testing and capacity building resulted in the development of the ARIES workspace with the following declared goals [15]:

**Figure 1.**
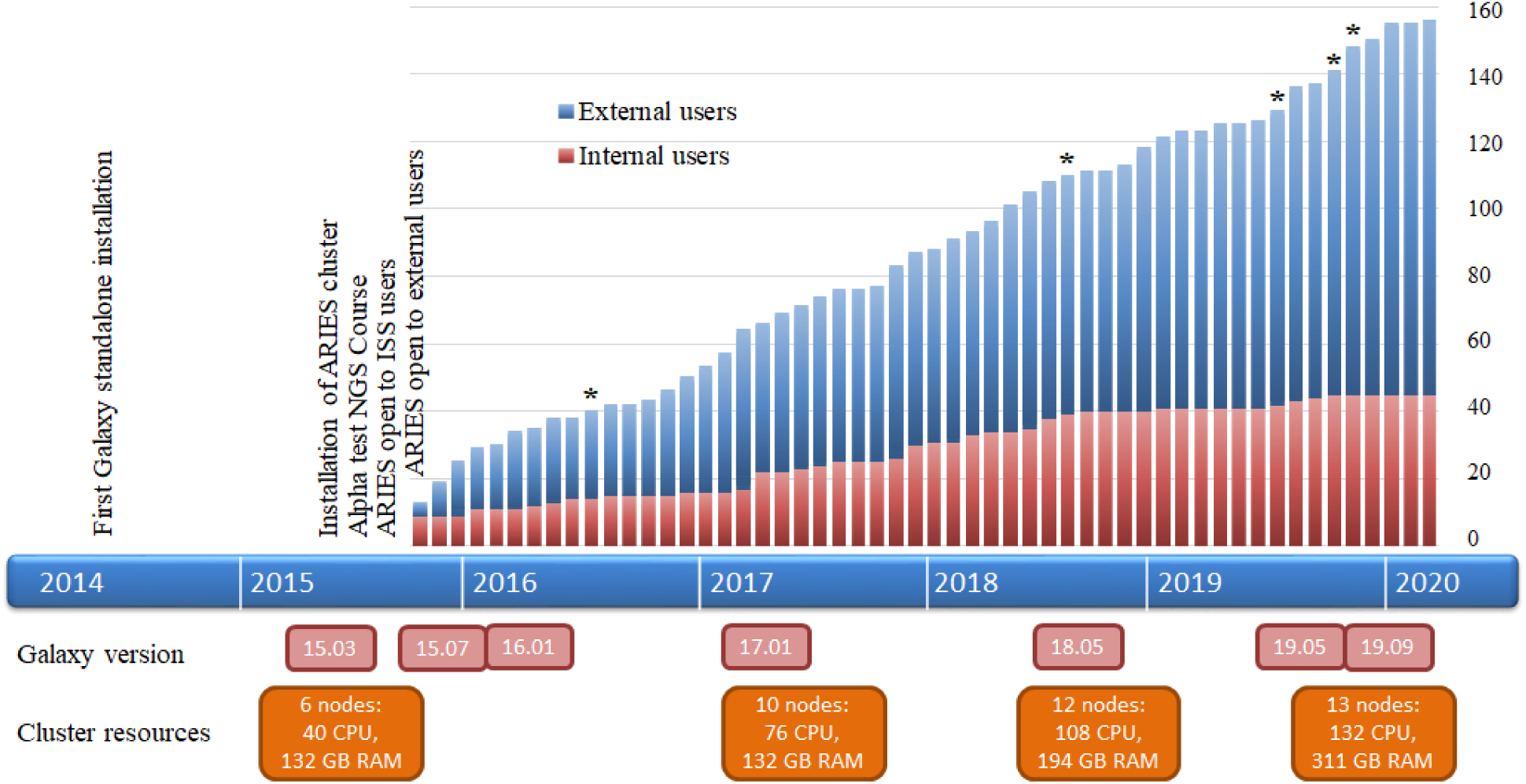
Timeline of the ARIES cluster experience. The number of internal and external users over time are shown. Asterisks (*) indicate when EURL NGS training courses were held utilising the ARIES platform. Underneath the timeline software and hardware upgrades are shown.

- Supporting scientists in genomic investigation of microbial pathogens, by providing at the same time a unique pre-configured genomic analyses environment characterised by reproducibility, transparency and easy data sharing as well as an open scalable platform for implementation of new bioinformatic resources.
- Development of an Information System for the collection of genomic and epidemiological data to enable the Next Generation Sequencing (NGS)-based surveillance of infectious diseases, foodborne outbreaks and diseases at the animal-human interface.
- Development of analytical pipelines enabling harmonised, real time multi-genome comparisons, to improve the detection of clusters of cases of infections and allowing the global bio-tracing of pathogens.
- Development of metagenomics models for the culture-independent detection and typing of pathogens and the study of their interactions with the microbiota in human and animal samples and in the vehicles of infections.

ARIES was installed in May 2015 (Figure 1) as a Galaxy instance for a multi-user production environment configured to run tools on a SLURM [16] cluster (https://galaxyproject.org/use/aries/). After a relatively short period of internal testing, an alpha-test was performed in June 2015 during a course on the bioinformatic analysis of NGS data organised by the EURL VTEC for the NRLs in the EU. The medium-small installation (at the time made up of 6 cluster nodes for a total of 40 CPU threads and 132 GB of RAM), could cope with the execution of jobs contemporarily launched by 16 users boosting confidence on the robustness of the infrastructure underlying the framework. Thus, in July 2015 ARIES was opened to internal users of the ISS and in October of the same year also to external users.

The number of users has steadily grown ever since and reached 156 accounts in March 2020, 45 of whom are scientists working at the ISS while 111 are external users (Figure 1, Table 1). Because of the mission of the EURL VTEC and the use of ARIES in six network training courses, many users are Europe-based NRL partners, however ARIES has also reached scientists outside this network and even in other continents. Internal users not only include personnel from the EURL (12 users) but also from other ISS Departments with interest in NGS applications different from public health microbiology. Moreover, users include drylab as well as wetlab scientists with or without background in bioinformatics.

**Table 1.**
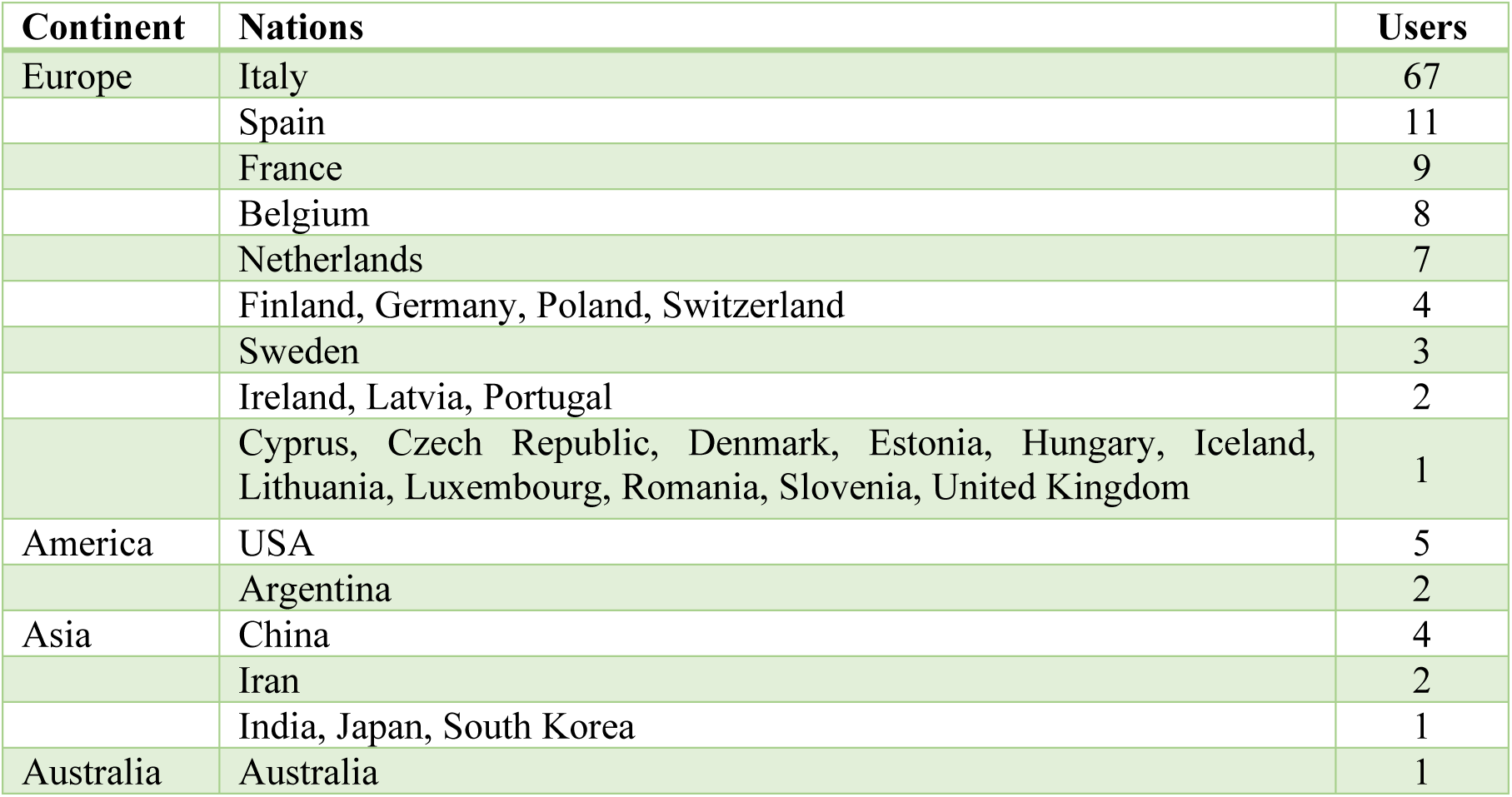
The number of user accounts per nation with access to ARIES in March 2020.

Most of the tools available on ARIES are dedicated to the analysis of bacterial genomes and particularly to *E. coli* WGS analysis. Nevertheless, specific databases for the analysis of genomes of other bacterial species are also available for specific tools. In detail, classical MLST typing is available for WGS of *Salmonella enterica, Listeria monocytogenes, Campylobacter jejuni, Clostridium botulinum, Clostridium difficile* and *Streptococcus pneumonia*. Additionally, cgMLST can also be performed for WGS of *Clostridium difficile, Legionella pneumophila, Listeria monocytogenes, Salmonella enterica* and *Staphylococcus aureus*. Tools dedicated to the analysis of viral sequences and for metagenomics are also available. More details are listed in the Methods section.

During these five years, ARIES has proven to be very stable with limited downtime, mainly for software upgrades. Hardware upgrades could almost always be implemented without interruption due to the cluster setup: only a migration of the underlying storage hardware required a system shutdown. Also because of the platform’s reliability, interest in the use of ARIES has continued to grow steadily over time and has not been limited to the initial enthusiasm (Figure 2).

**Figure 2.**
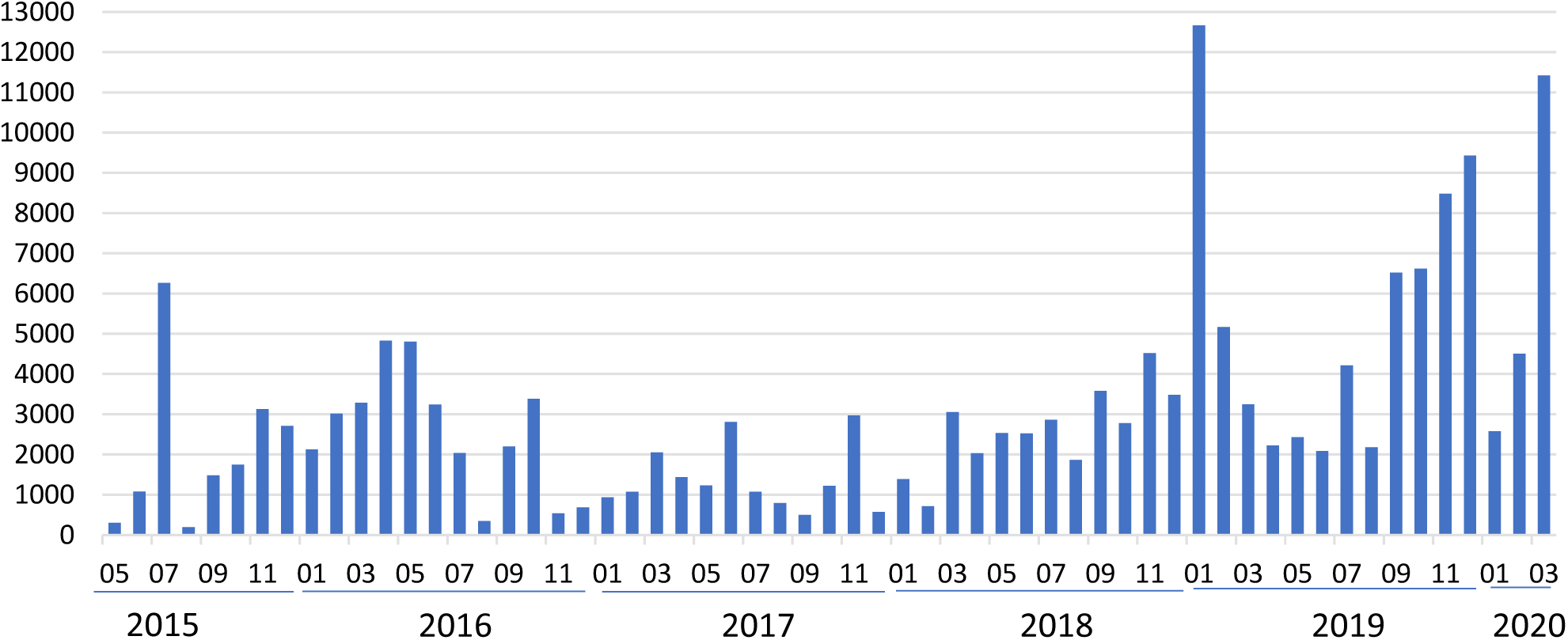
Historical chart of the number of jobs run per month on ARIES.

### EURL VTEC WGS Pipeline Tool

The EURL VTEC WGS PT (Pipeline Tool) was first developed in April 2017 as an effort to produce a simple typing tool for the analysis of *E. coli* short read sequencing data intended for non-bioinformatic users. This preconfigured workflow takes single or paired-end fastqsanger files as an input and performs a quality check on the sequences with FastQC [17] followed by trimming, assembly and various typing tools (Figure 3), resulting in an html summary report (Figure 4). From its first release, the pipeline has undergone several improvements, for instance by addition of Shiga toxin subtyping [18] and Antimicrobial Resistance (AMR) typing. A list of the released versions with an explanation of the main changes between them is available in the supplemental file S1_EURL_VTEC_WGS_PT.pdf.

**Figure 3.**
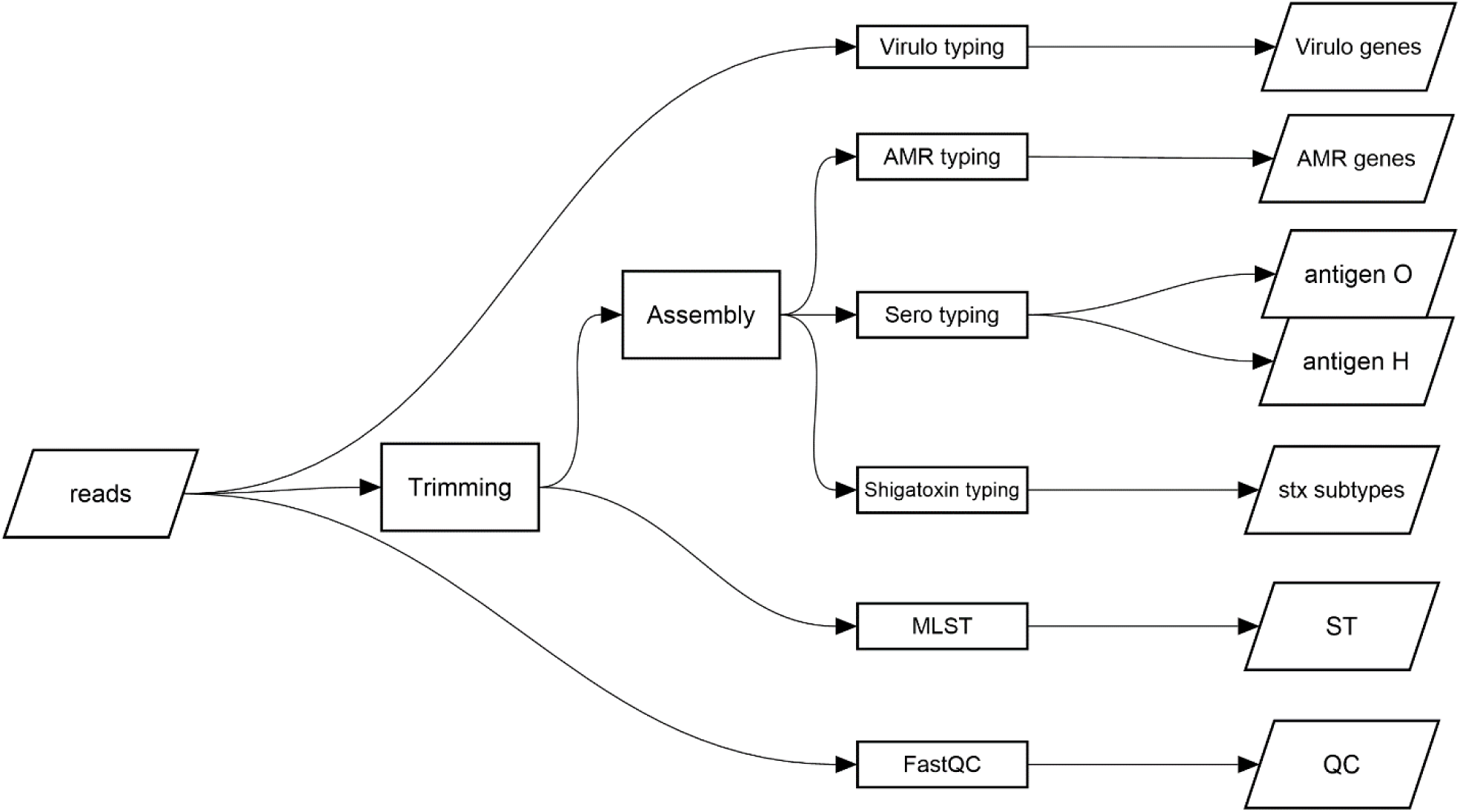
High-level flowchart of the EURL VTEC WGS version 3.1 pipeline tool. FastQC and Virulo typing are applied directly to the raw reads, while MLST analyses are performed with MentaLiST after trimming. Several typing tools are launched upon the assembled contigs. All the results are collected into an analysis report.

**Figure 4.**
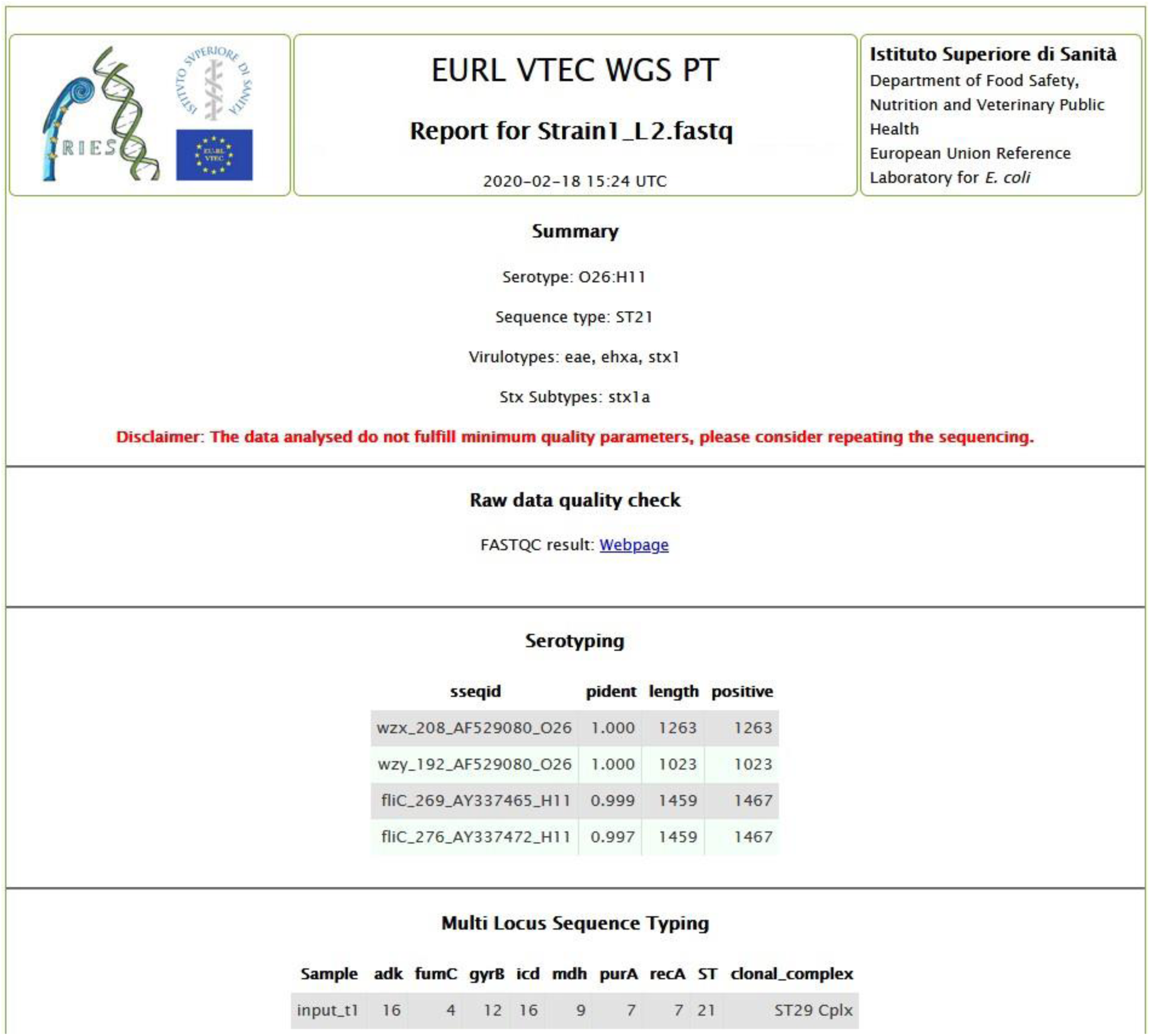
Example output report of the EURL VTEC WGS Pipeline Tool, the results of the typing tools are lined up in the summary section with detailed information attached below. A disclaimer warns that repeating sequencing should be considered if the data analysed does not meet the minimum requirement that the seven housekeeping genes of Multi Locus Sequence Typing are found and have at least a mean coverage of 30×.

In the current version 3.1, trimming is performed using fastq_positional_quality_trimming [19], assembly using SPAdes 3.12 [20] for single reads and INNUca 1.0 [21] for paired-end reads while MLST typing is done through MentaLiST 0.2.4 [8] using the seven housekeeping genes of the Achtman scheme for *E. coli* (adk, fumC, gyrB, icd, mdh, purA, recA) from Enterobase [22]. For AMR typing, the NCBI Antimicrobial Resistance Gene Finder [23] (AMRFinderPlus 3.6.7) is used. Serotyping of O and H antigens as well as Shiga toxin subtyping is obtained applying MMseqs2 to specific databases [24] (see the section Availability of supporting data). Virulence typing is based upon the patho_typer tool 1.0 [25] applied to the virulencefinder database from the Center for Genomic Epidemiology [26].

In the final comprehensive report of the pipeline (Figure 4), the results of the applied typing tools are lined up in a summary section, moreover detailed information is inserted in specific sections below.

Eventually, a disclaimer warns that repeating of sequencing should be considered if the data analysed does not meet the requirement that the seven housekeeping genes of Multi Locus Sequence Typing are found and have at least a mean coverage of 30×. The assembled contigs, a log file with the versions of tools and databases, the FastQC [17] and QUAST [27] html files and the typing output files are all available in the History menu.

### IRIDA-ARIES

From the experience of the EURL VTEC WGS pipeline, a further step to achieve the development of an Information System for the collection of genomic and epidemiological data to enable the WGS-based surveillance was made. To this means, an instance of the Integrated Rapid Infectious Disease Analysis (IRIDA) platform [28] that had been installed in November 2016, was personalised to implement an adapted version of this pipeline in August 2018 using ARIES as a workflow engine. The integrated IRIDA-ARIES platform is hosting the Italian national surveillance system for infections by *Listeria monocytogenes* (currently 551 samples) as well as the surveillance system for infections by *E. coli* (currently 626 samples, 338 human and 288 non-human). In this scenario, IRIDA communicates with ARIES through the latter’s unified Applications Programming Interface (API), completely masking the ARIES platform from the user who only interacts with the IRIDA user interface (Figure 5).

**Figure 5.**
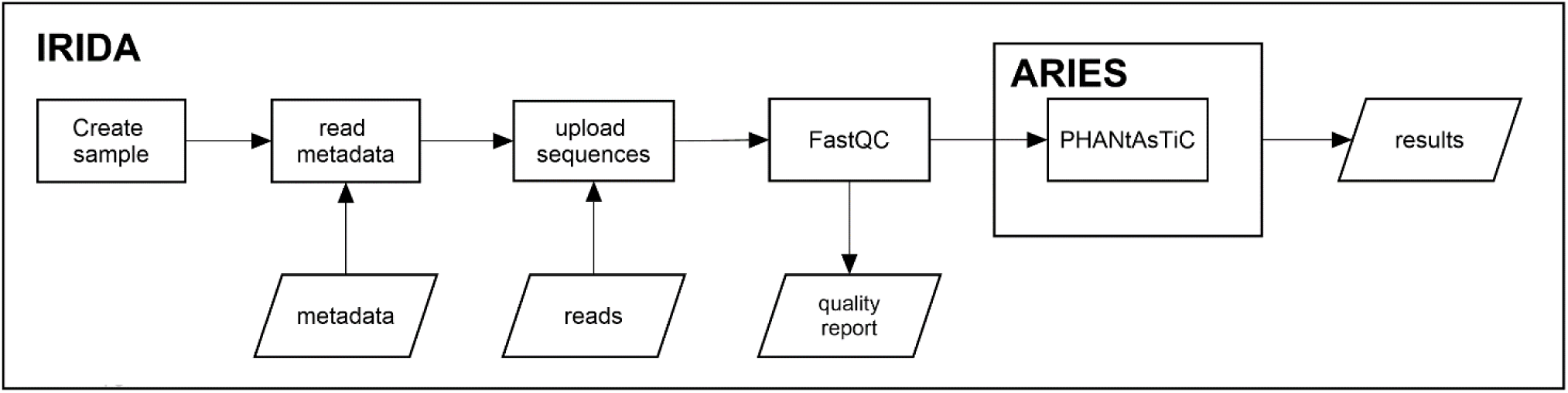
The integrated IRIDA-ARIES platform: when a sample is created metadata is recovered from the LIMS system. Then, when sequences are uploaded, the FastQC tool is applied and consequently the sequences are analysed in ARIES through the PHANtAsTiC pipeline. Results are returned to IRIDA and displayed in a specific analysis web page.

Whenever the user uploads a genomic sequence as a sample to a project, the automated analysis pipeline PHANtAsTiC 1.0 (Public Health Analysis of Nucleotides through Assembly, Typing and Clustering, Figure 6) is triggered, performing appropriate trimming (Trimmomatic [29]) and assembly (SPAdes or INNUca as before), typing and clustering elaborations in basis of the type of file (single-end reads versus paired-end reads) and bacterial species (*E. coli* or *Listeria*). The pipeline is executed on the ARIES cluster through an API call, using directly the sequences uploaded in IRIDA taking advantage of the shared file system and, upon completion, the results are sent back to IRIDA. All typing tools for *E. coli* are the same as those used in the EURL VTEC WGS pipeline (see specific section) while for *Listeria* the software LisSero 0.1 [30] determines this species’ serogroup. Quality assessment of the obtained assembly is performed by Quast 5.0.2 [27]. Cluster analysis applies cgMLST through chewBBACA 2.0.13 [9] for *E. coli* (INNUENDO scheme with 2360 loci [31]) and through MentaLiST 0.2.4 [8] for *Listeria* samples (Pasteur scheme with 1748 loci [32]). In both cases a distance matrix is calculated through a pairwise comparison between samples calculating their number of discordant alleles and a phylogenetic tree is built with the Bio.Phylo.TreeConstruction class of the BioPython package.

**Figure 6.**
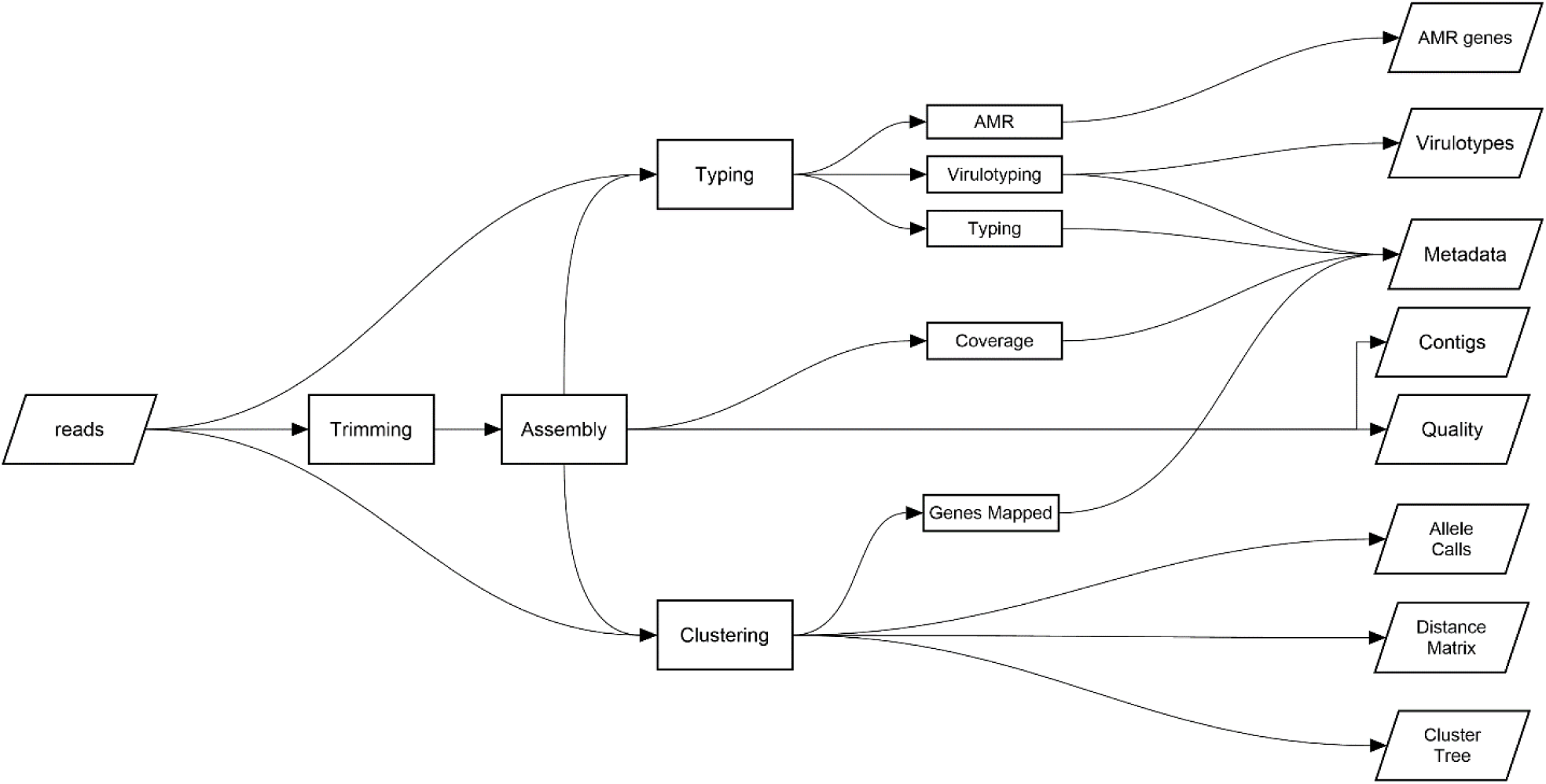
High-level flowchart of the PHANtAsTiC version 1.0 pipeline.

Currently IRIDA-ARIES has 60 users, predominantly personnel of the Regional Public Health Services (Public Health is delegated at the Regional level in Italy). ARIES is encapsulated with respect to IRIDA and able to automatically manage the input from the IRIDA platform without the need for direct user access, indeed only 15 IRIDA users also have ARIES accounts. Managers can access the Galaxy interface for debugging purposes in case of failures in the analysis process.

## Infrastructure experiences

The hardware components of the ARIES infrastructure are outlined in the Methods section below. For the current workload this system setup has proven to be stable and responsive. For instance, the PHANtAsTiC pipeline takes generally less than an hour to complete. Fibre channel storage was used for the cluster nodes’ operating systems, while a multiaccess Network File System (NFS) accommodates the shared file system that Galaxy demands for the cluster installation. The program autofs was applied for automatically mounting the shared file system on an as-needed basis and unmounting after a period of inactivity, avoiding any problem of freezing and unresponsivity of the file system. The new feature introduced in version 17.01 of the Galaxy platform enabling to install tool dependencies from Conda channels, allowed for tools published in the Galaxy ToolShed [12] to be installed flawlessly from the graphic user interface solving the infrastructures’ previous necessity of having to force permissions from the command line.

The communication between IRIDA and ARIES generally works fine: upon uploading and quality analysis of a sequence in IRIDA, the PHANtAsTiC pipeline gets fired off on ARIES and IRIDA polls the job until it finishes and then obtains the results in order to elaborate them into the samples metadata and produce the result page.

## Conclusions

Overall, the experience of using Galaxy as a platform for ARIES has been very positive in these five years. Currently, eleven journal articles and one report are listed as ARIES tagged publications [33] in the Galaxy Publication library [34]. The platform has proven extremely suitable for training and capacity building in the EURL VTEC network and is used by several NRLs as their primary WGS analysis tool. Thus, even with modest resources it has proven possible to obtain a stable and at the same time flexible platform, properties of great importance to the ARIES reality since the infrastructure in ISS is managed and maintained by one person only. A major improvement during the various versions of Galaxy has been the shift from ToolShed package recipes to Conda as the package management solution for the installation of software dependencies facilitating notably the integration of tools into the platform. The Galaxy framework provides users a visual tool for designing personalised workflows by themselves. This results very useful in the phases of debugging, fine-tuning and optimising analysis pipelines, while keeping track of all results as well as all settings including parameters for reproducibility. Importantly, data silos are avoided because it is simple to exchange analyses not only between users of ARIES but also with users on other Galaxy platforms. Easy collaboration is also possible for tools and pipelines, for example the EURL VTEC WGS PT has been shared with the United States Food and Drug Administration and is published since November 2019 on their GalaxyTrakr tool shed [35]. A solution for the shared NFS file system which represents an obstacle to the ability of the system to scale out is currently under investigation. Alternative suitable scenarios are the implementation of enterprise-level storage or migration of the cluster on a cloud infrastructure like the European Open Science Cloud (EOSC) [36] allowing for further scalability, fundamental for the platforms to retain responsiveness under intensive use, for instance in case of an outbreak investigation.

## Methods

### Main tools available in ARIES

General purpose tools for data upload/download/manipulation [12]

NGS Trimming: Trimmomatic [29], FASTQ positional and quality trimming [19]

NGS Assembly: SPAdes [20], SKESA [37], metaSPAdes [38], A5, INNUca [21], Shovill [39]

NGS Mapping: BWA [40], Bowtie2 [41]

NGS Alignment: BLAST [42], MMseqs2 [43], Diamond [44], MAFFT [45], MUMmer [46]

NGS Quality Control: FastQC [17], QUAST [27]

Phylogeny tools: PopPUNK [47], kSNP3 [48], FDA SNP Pipeline [49], MrBayes [50], PhyML [51], IQ-TREE [52]

Allele Call tools: SRST2 [53], MentaLiST [8], MLST [54], chewBBACA [9]

General Typing tools: Virulotyper [25], AMRFinderPlus [23], Resistance Gene Identifier [55]

Specific Typing tools: EURL VTEC WGS PT (*E. coli*), E coli Serotyper, E coli Virulo typer, E coli Shiga toxin typer, LisSero [30] (*Listeria*), SeqSero2 [56] (*Salmonella*)

Viral tools: Trinity [57], iVar [58], quasitools [59]

Metagenomics: Qiime2 [60], PICRUSt [61], ComMet [62], LefSe [63]

Visualisation: GraPhlAn [64], Krona [65], JBrowse [66]

### Infrastructure

The ARIES and IRIDA platforms have been set up mostly with hardware dismissed from institutional use, in fact no hardware has been acquired dedicated to this project except twenty hard disks for the QNAP Network Attached Storage (NAS).

Computation: Two HPE BladeSystem c7000 with each four GbE2c Layer 2/3 Ethernet Blade Switches and two HPE B-series 8/24c SAN Switches, dislocated in two different buildings of the Institute, constitute the core of the ARIES infrastructure. Currently, eight HPE ProLiant BL460c G6 servers are dedicated to IRIDA and ARIES. Alongside, from the beginning of 2020, an HPE Proliant DL380 G5 server and an HPE Proliant Proliant DL360p G8 server contribute to the cluster. The BladeSystem Enclosure was installed in 2007, while most hardware dates from 2012 (the G6 servers).

Storage: EMC VNX 5300 NAS was connected by Fiber Channel for the virtual machines’ Operating Sytems, QNAP TS-EC1680U NAS with Solid-State Drive cache acceleration was connected as an NFS for data storage.

Software: VMWare vSphere 6.0, Debian 9, Galaxy 19.09, IRIDA personalised fork from 19.01, SLURM 16.05, PostGreSQL 9.6, MariaDB 10.1, nginx 1.12.

### Availability of supporting source code and requirements

Project name: EURL VTEC WGS PT

Project home page: https://github.com/aknijn/eurl_vtec_wgs_pt-galaxy Platform: Galaxy

Programming language: Python and Perl, xml wrappers

Other requirements: dependencies are managed by the conda package management system.

License: EURL VTEC WGS pipeline is published under the GNU General Public License v3.0. All software from external sources are published under their respective licenses.

Project name: IRIDA-ARIES

Project home page: https://github.com/aknijn/irida-aries

Platform: IRIDA, IRIDA-ARIES is a fork from the IRIDA platform version 19.01

Programming language: Java and javascript

Other requirements: all dependencies are managed by the Apache Maven tool.

License: IRIDA-ARIES is published under the Apache License 2.0. All software from external sources are published under their respective licenses.

Project name: PHANtAsTiC

Project home page: https://github.com/aknijn/phantastic-galaxy Platform: Galaxy

Programming language: Python and Perl, xml wrappers

Other requirements: dependencies are managed by the conda package management system.

License: The PHANtAsTiC pipeline is published under the GNU General Public License v3.0. All software from external sources are published under their respective licenses.

### Availability of supporting data

*Escherichia coli* Virulo typing database: https://bitbucket.org/genomicepidemiology/virulencefinder_db/src/master/virulence_ecoli.fsa

*Escherichia coli* O antigen Serotyping database: https://bitbucket.org/genomicepidemiology/serotypefinder_db/src/master/O_type.fsa

*Escherichia coli* H antigen Serotyping database: https://bitbucket.org/genomicepidemiology/serotypefinder_db/src/master/H_type.fsa

*Escherichia coli* Achtman scheme MLST database: http://enterobase.warwick.ac.uk/species/ecoli/download_data

## Declarations

### Consent for publication

Not applicable.

### Competing interests

The authors declare that they have no competing interests.

### Funding

This work is supported by the ISS intramural funds.

### Authors’ contributions

S.M. ideated and coordinated the projects of ARIES and IRIDA. A.K. planned and performed the installation of ARIES and IRIDA, A.K. adapted the IRIDA software. M.O. and A.K. coded bioinformatic tools for ARIES. All authors participated in development of bioinformatic tools and analysis pipelines. A.K. coordinated the writing process. All authors contributed to the writing process and to the critical review of the manuscript.

## List of abbreviations

AMR: Antimicrobial Resistance
API: Application Programming Interface
ARIES: Advanced Research Infrastructure for Experimentation in genomicS
CPU: Central Processing Unit
ECDC: European Centre for Disease Prevention and Control
EFSA: European Food Safety Authority
*E. coli*: *Escherichia coli*
EURL: European Union Reference Laboratory
GB: GigaByte
IRIDA: Integrated Rapid Infectious Disease Analysis
ISS: Istituto Superiore di Sanità
MLST: MultiLocus Sequence Typing
NAS: Network Attached Storage
NFS: Network File System
GS: Next Generation Sequencing;
NRL: National Reference Laboratory
PFGE: Pulsed-field Gel Electrophoresis
PHANtAsTiC: Public Health Analysis of Nucleotides through Assembly, Typing and Clustering
PT: Pipeline Tool
RAM: Random Access Memory
VTEC: Verotoxigenic *Escherichia coli*
WGS: Whole Genome Sequencing

## Acknowledgements

The authors thank Amalia Ferretti for designing the ARIES logo.

## Notes

https://aries.iss.it

### Competing Interest Statement

The authors have declared no competing interest.

