## Supplemental Table S1 for "Advanced Research Infrastructure for Experimentation in genomicS (ARIES): a lustrum of Galaxy experience"

### Additional files

Release dates of the developed versions of the EURL VTEC WGS Pipeline Tool with an explanation of the main changes between them indicated in the last column.

| V | Release Date | Changes |
|---|---|---|
| 0.1 | April 2017 | First version with all tools locally copied in specific folders (A5, BLAST, Bowtie2, FastQC, NGS_Quality, SequenceTyper, SPAdes, ViruloTyper) |
| 1.0 | December 2017 | Version with a first attempt to delegate dependencies to Conda; the ViruloTyper tool has been substituted by the INNUENDO Patho_Typer tool (with personalised virulence database). |
| 1.1 | May 2018 | The dependencies are now installed through ARIES by Conda. The xml wrapper file has been validated by planemo, test data has been added. The pipeline outputs a log file explicating the versions of the tools and databases used in the anlysis. |
| 1.2 | June 2018 | The virulotyping and serotyping steps can now be included or excluded from the analysis. |
| 2.0 | June 2018 | Only SPAdes is used for assembly. For serotyping, before a filter step is introduced applying duk. For MLST of paired-end reads, the two files are put together with the input_se (works better). The fastq filtering step can now be included or excluded from the analysis. |
| 2.1 | August 2018 | Possibility to use a paired-end collection of fastqsanger files as an input. |
| 2.2 | October 2018 | Shiga toxin typing added, the step can be included or excluded from the analysis. |
| 2.3 | September 2019 | SRST2 is substituted by MentaLiST for Multi Locus Sequence Typing. |
| 3.0 | January 2020 | The duk filter steps are removed, a total assembly is performed using SPAdes. BLAST is substituted by mmseqs2 in serotyping and Shiga toxin typing. AMRtyping is performed by the AMRFinderPlus tool. |
| 3.1 | April 2020 | The Shiga toxin typing tool has been modified thoroughly to improve effectiveness. A consensus is created through combined assembly and mapping steps and a BLAST search is run on the result. |
